# Functional connectivity is dominated by aperiodic, rather than oscillatory, coupling

**DOI:** 10.1101/2024.09.18.613682

**Authors:** N. Monchy, J. Duprez, J-F. Houvenaghel, A. Legros, B. Voytek, J. Modolo

## Abstract

Functional connectivity has attracted significant interest in the identification of specific circuits underlying brain (dys-)function. Classical analyses to estimate functional connectivity (i.e., filtering electrophysiological signals in canonical frequency bands and using connectivity metrics) assume that these reflect oscillatory networks. However, this approach conflates non-oscillatory, aperiodic neural activity with oscillations; raising the possibility that these functional networks may reflect aperiodic rather than oscillatory activity. Here, we provide the first study quantifying, in two different human electroencephalography (EEG) databases, the contribution of aperiodic activity on reconstructed oscillatory functional networks in resting state. We found that more than 99% of delta, theta, and gamma functional networks, more than 90% of beta functional networks and between 23 and 55% of alpha functional networks were actually driven by aperiodic activity. While there is no universal consensus on how to identify and quantify neural oscillations, our results demonstrate that oscillatory functional networks are drastically sparser than commonly assumed. These findings suggest that most functional connectivity studies focusing on resting state actually reflect aperiodic networks instead of oscillations-based networks. We highly recommend that oscillatory network analyses first check the presence of aperiodicity-unbiased neural oscillations before estimating their statistical coupling to strengthen the robustness, interpretability, and reproducibility of functional connectivity studies.

## INTRODUCTION

Functional connectivity (FC) is a staple in cognitive neuroscience. The identification of specific brain networks provides insights into (patho-) physiological processes (Medaglia et al., 2015). In both electroencephalography (EEG) and magnetoencephalography (MEG), FC consists in identifying the statistical couplings of neural oscillations from different brain regions of interest (ROIs; Friston et al., 1993). Neural oscillations are usually defined on specific frequency bands as delta (1–4 Hz), theta (4–8 Hz), alpha (8–12 Hz), beta (12–30 Hz), gamma (30-140Hz).

FC can be estimated using a plethora of metrics, such as amplitude- (e.g., Amplitude Envelope Correlation, AEC) or phase-based (e.g., Phase Locking Value, PLV) measures, which all have associated assumptions, such as ignoring zero-lag connectivity as with Phase Lag Index (PLI). Despite extensive studies aiming at evaluating the conditions in which FC metrics are optimal, there is not a clear consensus on FC metrics that would provide a reliable estimate of dynamic brain connectivity under all circumstances (Fraschini et al., 2020; Allouch et al., 2023).

Classical analyses to estimate FC involved the following : acquisition of physiological signals, preprocessing, filtering of clean signals within a specific frequency band of interest for each ROI, and then computing a FC metric value for each possible pair of signals, resulting in a *N* x *N* symmetrical matrix (with zeros on the diagonal, since self-connections cannot be evaluated) that provides the “strength” of statistical coupling between the *N* ROIs. These usual analyses make the assumption that it reflects oscillatory networks.

However, as pointed by Donoghue et al., a major bias in the estimation of oscillatory power from spectra lies within the presence of the so-called “aperiodic” activity (Donoghue et al., 2020). Aperiodic activity exhibits a 1/f-like distribution, with power decreasing exponentially across increasing frequencies. This component, often described by a 1/*f* function, is parameterized by two parameters : the exponent reflecting the aperiodic power pattern across frequencies and the offset indicating a uniform power shift across frequencies.

To our knowledge, the quantitative contribution of aperiodic activity in the estimation of functional networks has not yet been studied. We hypothesized that aperiodic activity had a great quantitative contribution in the estimation of functional networks. To address this, we analyzed whether and how aperiodic activity impacts the reconstruction of oscillatory functional networks in two different human EEG databases. We followed the usual analysis pipeline and compared it to a new approach verifying the presence of oscillations not conflated by aperiodic activity, referred to as ‘aperiodicity-unbiased oscillations’. We chose to analyze resting-state recordings due to their consistent brain connectivity patterns across subjects and methods (Brookes et al., 2011; Damoiseaux et al., 2006) which are essential for both cognitive (Alavash et al., 2015) and clinical studies (Hassan et al., 2017).

## MATERIAL AND METHODS

In this study, we applied the same analysis pipeline to two independent datasets (Figure 1).

**Figure 1.**
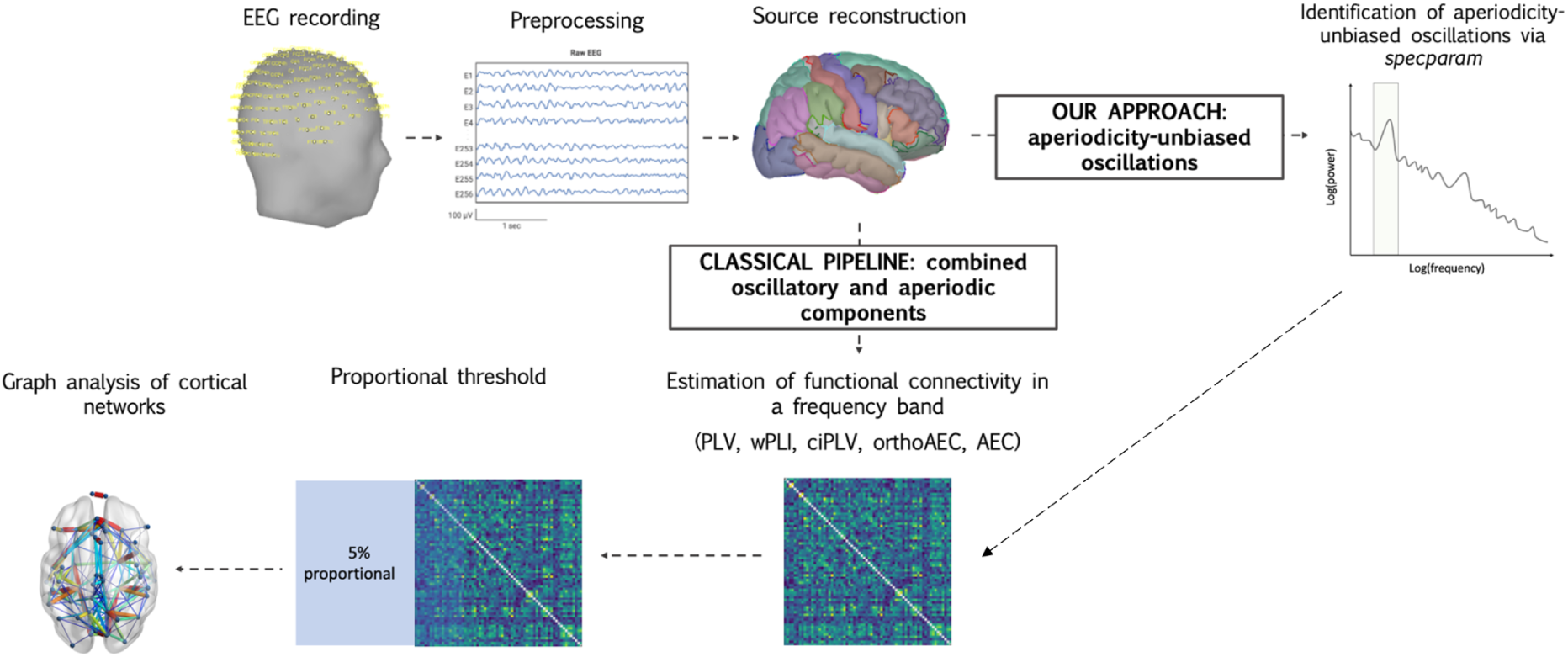
Analysis pipeline of the study. After EEG recordings, signals were preprocessed to remove artifacts. The EEG inverse problem was solved using the weighted minimum norm estimate method and cortical sources were projected on the Desikan-Killiany Atlas. Then, two approaches were conducted: the classical pipeline and our approach that separately considers the periodic and aperiodic components of the signal. This approach permits identifying regions with aperiodicity-unbiased oscillations before the estimation of FC. After computing FC with 5 different metrics, a proportional threshold was applied to retain the top 5% of the strongest connections. Finally, the functional networks were characterized using graph metrics.

### Participants

#### Dataset A

Thirty HC (15 females) aged between 45-70 years old (mean = 61.7, sd = 7.3) and 29 patients (15 females) diagnosed with idiopathic PD (Hughes et al., 1992), aged between 45-73 years (mean = 60.4, sd = 7.3) participated in this study. HC and patients did not significantly differ in age, sex or education. All patients were recruited from the Neurology Department of Rennes University Hospital (France).

All participants were free from any major cognitive impairment (Montreal Cognitive Assessment Scale, MoCA scores < 22; Nasreddine et al., 2005) or severe neurocognitive disorder according to the Diagnostic and Statistical Manual of Mental Disorders - V (DSM-V). Participants with moderate or severe psychiatric symptoms and present or past neurological pathology (other than PD for patients) were not included in this study. For further details, see Monchy et al., 2024.

This study was conducted in accordance with the declaration of Helsinki and was approved by a national ethics committee (CPP ID-RCB: 2019-A00608-49; approval number: 19.03.08.63626). After a complete description of the study, all participants gave their informed written consent.

#### Dataset B

One hundred and three HC (62 females, 41 males) aged between 17-71 years old (mean = 37.9, sd = 13.9) who participated in an experimental paradigm used in the *Stimulus-Selective Response Modulation* (SRM) project at the Dept. of Psychology (University of Oslo, Norway) were studied (Hatlestad-Hall et al., 2022). None of the subjects reported severe psychiatric or neurological symptoms. For further details, see Hatlestad-Hall et al., 2022.

### HD-EEG recording and processing

#### Recording

##### Dataset A

EEG data was recorded using a High-Definition EEG (HD-EEG) system (EGI, Electrical Geodesic Inc., 256 channels) with a sampling frequency of 1000 Hz. Electrode impedance was maintained below 25 kΩ. The Cz electrode was used as reference, and the ground close to Pz. Electrodes on the net were placed using the standard 10-10 geodesic montage. Most jaw and neck electrodes were removed due to excessive muscular artifacts, resulting in a total of 199 exploitable electrodes, as shown in the channel file available in the GitHub repository (see Section Data and code availability). The EEG recording consisted of a 5-minutes resting state period with eyes closed.

##### Dataset B

EEG signals were recorded from a 64-channel (Ag-AgCl electrodes) BioSemi ActiveTwo system. Electrodes were positioned according to the extended 10-20 system (10-10). Data were sampled at 1024 Hz without online filters, except for the default hardware anti-aliasing filter. EEG data was recorded during 4 minutes in resting-state with eyes closed.

#### Preprocessing

##### Dataset A

All EEG preprocessing and subsequent analyses were performed manually using the Brainstorm toolbox (Tadel et al., 2011) in Matlab (MathWorks^®^, USA). First, DC offset removal was applied. Second, a notch filter (50 Hz) and a band-pass filter (1-90 Hz) were applied (Finite Impulse Response; FIR filter using Kaiser window). Third, signals were visually inspected and bad channels were removed before being interpolated using Brainstorm’s default parameters. Fourth, Independent Component Analysis (ICA, *jade* method) was used to remove artifacts following visual inspection of the independent components. Fifth, signals were segmented into 4-second epochs. Finally, a visual inspection was performed to manually reject remaining bad epochs. As a result, an average of 59 (sd = 14) resting-state epochs were studied per subject.

##### Dataset B

Preprocessing of EEG data has already been achieved upon download. EEG data was preprocessed using a basic and fully automated pipeline on EEGLAB (Delorme & Makeig, 2004) and NoiseTools (De Cheveigné & Arzounian, 2018) in Matlab (MathWorks^®^, USA). First, the pipeline identified and removed bad channels. Second, a high-pass filter of 1 Hz was applied. Third, power line noise (50 Hz) was removed with Zapline (De Cheveigné, 2020). Fourth, ICA was calculated using the SOBI algorithm (Belouchrani et al., 1997). All components of ocular or muscular origin with a certainty above 85% were subtracted from data. Fifth, removed channels were interpolated. Sixth, a low-pass filter of 45 Hz was applied. Sixth, bad channels characterized by repeating small signal glitches were removed. Finally, data were referenced to the average reference and segmented into non-overlapping 4-second epochs. Epochs containing discontinuities were removed. This included fully automated cleaning procedure, considered strict in terms of the data rejection, produced a derived dataset in which 64.1% of epoched data files kept more than 90% of their channels and 23.5% retained between 75% and 90% of their channels.

#### Source reconstruction

We used a realistic head model and electrode position. Here, we used the Boundary Element Method (BEM) head model fitted to the International Consortium for Brain Mapping Magnetic Resonance Imaging (ICBM MRI) template (Mazziotta et al., 2001), and the OpenMEEG toolbox (Gramfort et al., 2010). The EEG inverse problem was solved using the weighted minimum norm estimate (wMNE) method (Lin et al., 2006). Cortical sources were projected on the Desikan-Killiany Atlas parcellation (68 regions of interest; ROIs) (Destrieux et al., 2010), with their orientation constrained normally to the cortical surface (Dale & Sereno, 1993). A noise covariance matrix was computed from our 700 ms pre-stimulus baseline. Signal-to-noise ratio (SNR) and depth weighting were set to default Brainstorm values.

### Spectral parameterization

In our approach, we specifically parameterized the EEG signal into both aperiodic activity and oscillations in order to identify oscillations that are not conflated by aperiodic activity. For each subject, frequency band and ROI, we checked whether ‘true’ oscillations, not conflated by aperiodic activity, were found by *specparam*. In this study, we used the term ‘aperiodicity-unbiased oscillations’ to refer to oscillations not conflated by aperiodic activity according to the *specparam* algorithm.

Power spectrum density (PSD) was computed for each epoch using Welch’s method (window length of 0.5 second, 50% window overlap), which is a necessary step for subsequent estimation of aperiodic parameters. Spectra were then averaged by subject. For each resulting power spectrum, we applied the spectral parameterization algorithm in the 1–40 Hz frequency band (*specparam* toolbox version 1.0; Donoghue et al., 2020), which considers the PSD as a linear combination of two different types of components: aperiodic and periodic (oscillatory) components (Donoghue et al., 2020). The aperiodic component is estimated through aperiodic exponent and offset while the periodic component is assessed with the frequency and amplitude of oscillatory peaks. In order to tailor the *specparam* algorithm to our data, we compared the goodness-of-fit metrics of several models, ultimately selecting the one with optimal fitting metrics: (dataset A : channel-averaged R^2^ = 0.88; channel-averaged mean squared error, MSE = 0.02; dataset B : channel-averaged R^2^ = 0.9; channel-averaged mean squared error, MSE = 0.04). Optimized parameters included: peak width limits: [0.5-6]; max number of peaks: 4; minimum peak height: 1.0; peak threshold: 2.0 and aperiodic mode = ‘fixed’. Aperiodic parameters were estimated at the ROI level.

### Functional connectivity

Numerous approaches have been proposed for calculating FC between reconstructed regional time series. Here, we chose to compute different widely-used EEG functional connectivity estimation methods based on phase or amplitude. For each subject, each frequency band, and each epoch, FC was calculated using these different methods to obtain a 68x68 connectivity matrix, where each entry *a*_*i*,*j*_ of the FC matrix represented the weight of the connection linking *ROI*_*i*_ to *ROI*_*j*_ . All FC matrices were computed on Brainstorm (Tadel et al., 2011).

#### Phase-based metrics

FC can be defined by the relative instantaneous phase between two time series. Phase-based metrics are robust against variations in signal amplitude, but low signal-to-noise values remain challenging for these measures (Lachaux et al., 1999; Mormann et al., 2000).

### i. Phase Locking Value (PLV)

The most commonly used phase measure is the Phase Locking Value (PLV), defined as the absolute value of the mean phase difference between two signals, expressed as a complex unit-length vector (Lachaux et al., 1999; Mormann et al., 2000). When the distribution of the phase difference between two signals is uniform, PLV is 0. Conversely, if the phase difference between the two signals has a preferred angle, PLV approaches 1.

Considering a pair of narrow-band analytic signals, *i* and *j*, PLV is defined as : 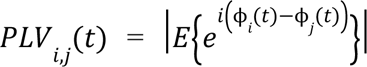 where *E*{.} denotes the expected value and ϕ(*t*) is the instantaneous phase derived from the Hilbert transform.

### ii. Corrected imaginary PLV (ciPLV)

PLV has a major limitation in assessing functional brain connectivity, namely its sensitivity to volume conduction (Stam et al., 2007) and source-leakage effects. In source-level EEG data, neighboring sources may still share some activity due to data low spatial resolution (source leakage). To face this limitation, Bruña and colleagues (Bruña et al., 2018) proposed phase-based metrics that discard zero-lag connectivity, thereby limiting volume conduction.

Considering a pair of narrow-band analytic signals, *i* and *j*, ciPLV is defined as: 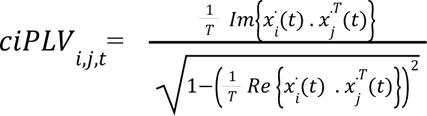 where *Im*{*x*} and *Re*{*x*} stands respectively for the imaginary and real part of *x*, 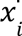 is the oscillatory part of analytical signal *i*, *x*^.*T*^ is the transpose conjugate of 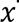 and T is the data length.

### iii. Weighted Phase Lag Index (wPLI)

Stam and colleagues also proposed the phase lag index (PLI) as another phase-based metric (Stam et al., 2007). PLI measures the asymmetry of the distribution of phase differences between two signals, and is an efficient method to detect “true” changes in phase-synchronization, and has a low sensitivity to volume-conducted noise. However, its sensitivity to noise and volume conduction could be impacted by its discontinuity, particularly affecting synchronization effects of minimal magnitude.

Considering a pair of narrow-band analytic signals, *i* and *j*, the complex cross-spectrum *C* for two signals is computed by Fourier-transforming them into *X*(*f*) and *Y*(*f*). *X* and *Y* are used to compute the cross-spectrum *C*(*f*)= *X*(*f*) *Y* (*f*), where *Y* indicates the complex conjugate of *Y*. If we focus on a specific frequency *f*\*, the complex non-diagonal part of *C* is called *Z*. ℑ corresponds to the imaginary part of Z. wPLI is defined as:

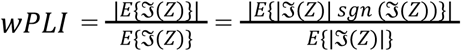

#### Amplitude-based metrics

Another coupling mode between neuronal oscillations from different brain regions operates purely on the amplitude envelope correlation, reflecting the temporal co-modulation of amplitude (Bruns et al., 2000; Siegel et al., 2012). Amplitude correlation is considered as an efficient index of the large-scale cortical interactions mediating cognition (Mazaheri, 2010).

### i. Amplitude Envelope Correlation (AEC)

Amplitude Envelope Correlation (AEC) is computed by correlating the amplitude envelopes of two oscillatory signals. This normalized measure of amplitude coupling is based on linear correlations between signals’ envelopes derived from the Hilbert transform (Brookes et al., 2011; Hipp et al., 2012). High AEC values (close to 1) correspond to synchronous amplitude envelope fluctuations between oscillations.

### ii. Orthogonalized Amplitude Envelope Correlation (orthoAEC)

This metric is a variant of AEC that removes zero-lag signal overlaps due to spatial leakage through a multivariate symmetric orthogonalization approach (Colclough et al., 2015). Signals are orthogonalized before derivation of their envelopes, and then the linear correlation between these envelopes are computed.

In the classical approach, FC estimation is performed directly after source reconstruction. In parallel with classical analyses, our approach estimated FC after identifying the brain regions that exhibit aperiodicity-unbiased oscillations. Each FC matrix coefficient *a*_*i*,*j*_ was retained only if an aperiodicity-unbiased oscillation between [1-45] Hz was reported by the specparam model in *ROI*_*i*_ and in *ROI*_*j*_, implying the presence of actual oscillations in two regions of interest.

#### Thresholding method

After estimating FC, we computed for each different FC method and for each frequency band, the epoch-averaged connectivity matrix for each subject. A usual step in network estimation is FC matrix thresholding, thereby removing spurious connections and resulting in sparsely connected matrices which are necessary for the computation of most graph theoretical metrics. This proportional threshold retains the strongest correlations of the connectivity matrix. This selection is often mentioned as an analysis in which the “density” (Jalili, 2016; Van Den Heuvel et al., 2008) or “network cost” (Achard & Bullmore, 2007; Bassett et al., 2008; Ginestet et al., 2011) is fixed across groups. Thresholding value choice is arbitrary. Here, according to standard literature values, we set the proportional threshold of connectivity matrices at 5%.

After thresholding connectivity matrices, subject-averaged FC matrices were calculated for each FC estimation method and for each frequency band.

### Comparison metrics

#### Descriptive aspects

After estimating cortical FC networks, the number of nodes and the percentage of connections were computed for each frequency band in both approaches to describe the architecture of networks.

#### Graph metrics

To describe the topological graph properties of each cortical network, we also calculated the graph metrics most commonly used in the literature (Rubinov & Sporns, 2010; De Vico Fallani et al., 2014). We chose widely-employed measures detecting functional integration and segregation, centrality, and testing network resilience taking into account their ability to characterize large (whole network), intermediate (sub-networks) or small (nodes) topological scales. Graph metrics were computed using the Brain Connectivity Toolbox (BCT; Rubinov et al., 2009).

### i. Clustering coefficient

The clustering coefficient is the fraction of triangles around a node which corresponds to the fraction of the node’s neighbors that are also neighbors of each other (Watts & Strogatz, 1998). This measure is indicative of the prevalence of clustered connectivity around individual nodes, and more globally of network segregation.

### ii. Characteristic path length

Paths are sequences of distinct linked nodes that never visit a single node more than once, and represent potential routes of information flow between brain regions. The length of these paths indicates the potential for functional integration between brain regions, shorter paths underlying stronger potential for integration. The average shortest path length between all pairs of nodes in the network is called characteristic path length (Watts & Strogatz, 1998), which is the most common measure of functional integration.

### iii. Global efficiency

Global efficiency corresponds to the average inverse shortest path length (Latora & Marchiori, 2001). This measure may be computed on disconnected networks, as paths between disconnected nodes are defined to have infinite length (zero efficiency). Global efficiency is often considered as a superior measure of integration (Achard & Bullmore, 2007).

### iv. Betweenness centrality

Betweenness centrality is the fraction of all shortest paths in the network that contain a given node. Nodes with high betweenness centrality values indicate participation in a large number of shortest paths. This centrality measure is based on the principle that central nodes participate in many short paths within a network, thereby acting as important centers of information flow (Freeman, 1977).

### v. Degree distribution

The degree of an individual node is the number of links connected to that node, corresponding to its number of neighbors. Nodes with high degree are defined as hubs. The degree distribution is the distribution of degrees of all network nodes (Barabasi & Albert, 1999). Degree values reflect the importance of nodes, while the degree distribution is a marker of network development and resilience. This indirect measure of resilience quantifies anatomical features reflecting network vulnerability to random node removal (attacks). Most real-life networks do not have a degree distribution that perfectly conforms to the power law, but rather behaves locally like a power law. The extent to which these distributions correspond to a power law is therefore a useful marker of resilience (Achard et al., 2006).

### Statistical analysis

All statistical analyses were conducted in R v.4.1.3. (R Core Team, 2023) implemented with the tidyverse and lme4 packages (Wickham et al., 2019; Bates et al., 2015). The significance threshold was set at 0.05.

In order to assess the quantitative contribution of aperiodic activity on the estimation of FC networks, we compared graph metrics values obtained from connectivity matrices derived from the classical pipeline *versus* those obtained from our approach identifying the aperiodicity-unbiased oscillations. Dataset A involved two population groups (HC and PD), we tested both the effect of group and approach using two-factors ANOVA with repeated measures. For each method of FC estimation and for each frequency band, an ANOVA model was performed as follows: anova_test(graph_metric, wid = subject, between = group, within = approach). Conditions of application were checked for each model. When sphericity assumption was not met, a Greenhouse-Geisser correction was applied. In the case of Dataset B, the effect of the selection of aperiodicity-unbiased oscillations was assessed with paired t-tests. If the data distribution shows a very large majority of values equal to 0, statistical tests have not been carried out (non-normal data). Moreover, in the presence of outliers, it was verified that the statistical tests remained significant when outliers were excluded from the test. False Discovery Rate (FDR) corrections p-values were computed following multiple comparisons.

Pearson correlations between FC matrices derived from the classical pipeline and FC matrices from our approach were computed, for each FC estimation method, as a global measure of similarity between alpha cortical networks. Due to the multiple comparisons between each FC method, an FDR correction was applied.

### Data and code availability

All Matlab and R codes used for processing and data analysis are publicly available at (https://github.com/noemiemonchy/1-f-corrected_functional_networks).

#### Dataset A

The channel file, and all Matlab and R codes used for preprocessing and data analysis are publicly available at (https://github.com/noemiemonchy/PD-SIMON).

#### Dataset B

SRM resting-state EEG data are publicly available (https://OpenNeuro.org).

## RESULTS

Since the results were similar across each FC method, only those obtained using the PLV and wPLI methods are presented for readability. The results from ciPLV, orthoAEC, and AEC methods are available in Supplementary Figures 1-3.

### Sparsity of functional networks computed on aperiodicity-unbiased oscillations

The quantitative contribution of aperiodic activity on FC estimation was evaluated by comparing, for each dataset, FC matrices from the classical pipeline, with FC matrices from our approach considering only aperiodicity-unbiased oscillations. (Figure 2).

**Figure 2.**
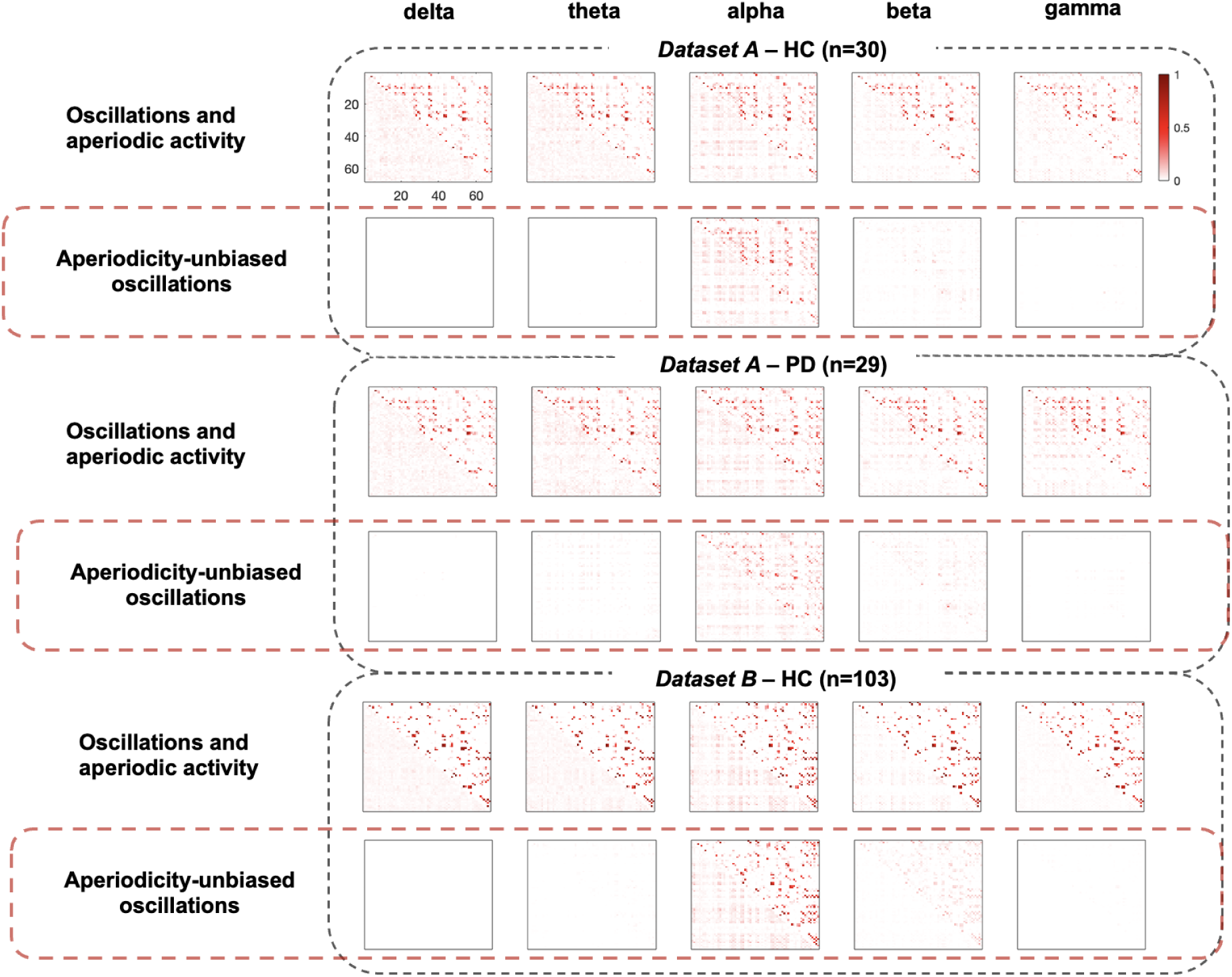
Functional connectivity matrices averaged across subjects for each dataset and each frequency band. The upper triangular portion of the matrices presents FC results from PLV, and from wPLI for the lower portion. Results are presented for each dataset: the first row shows FC matrices from the classical pipeline and the second row displays FC matrices from the approach verifying the presence of oscillations not conflated by aperiodic activity.

First, subject-averaged connectivity matrices derived from the classical pipeline exhibited a similar pattern between ROIs across all frequency bands for each method. Notably, FC values were higher in the alpha band. These results were consistent across all studied datasets. Additionally, FC was overall lower when looking at methods correcting for zero-lag.

FC matrices computed on aperiodicity-unbiased oscillations showed almost no connectivity in delta and gamma bands. Only the alpha band maintained FC values similar to matrices derived from the classical pipeline. A few FC values persisted in the theta and beta bands, however overall connectivity in these frequency bands was greatly reduced. The selection of aperiodicity-unbiased oscillations resulted in sparser connectivity matrices across frequency bands. These results were consistent across all studied datasets.

For all datasets, we studied the number of nodes and the number of connections that were maintained with the selection of aperiodicity-unbiased oscillations. The number of nodes was higher in the alpha band with an average of 47 nodes (sd = 11) and 39 nodes (sd = 25) for the A-HC and PD data respectively, and an average of 58 nodes (sd = 14) for the dataset B (Figure 3). In the beta band, some nodes were also found across all datasets : an average of 11 nodes (sd = 6) were kept in the dataset A-HC, approximately 13 nodes (sd = 8) in dataset A-PD and around 13 nodes (sd = 14) in dataset B. Nodes in delta, theta and gamma bands were much sparser. On average, in the theta band, no nodes (sd = 1) were kept in A-HC, about 3 nodes (sd = 6) were found in dataset A-PD and only 1 node (sd = 5) was maintained on dataset B. Finally, in the gamma band, around 1 node (all datasets : sd = 3) was found and while no node (dataset A-HC : sd = 0; dataset A-PD : sd = 1, dataset B : sd = 0) was found in the delta band for each dataset.

**Figure 3.**
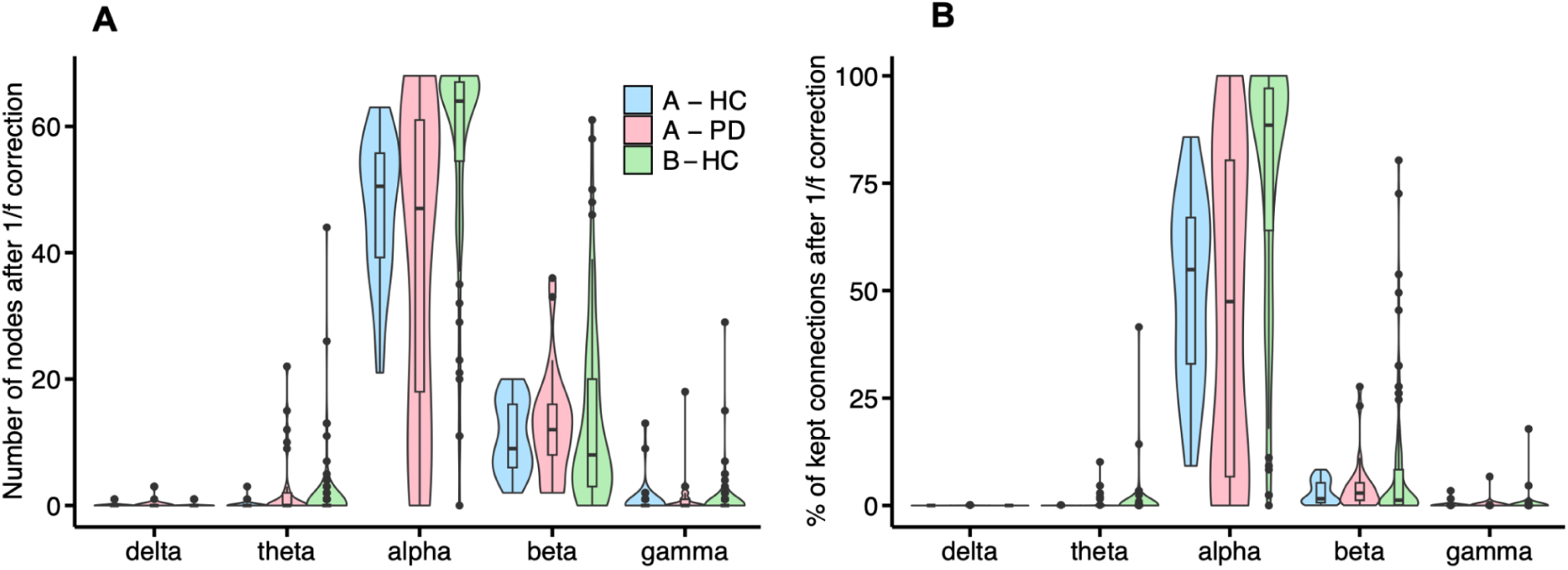
Number of nodes (A) and percentage of kept connections (B) after identifying the presence of aperiodicity-unbiased oscillations for each frequency and dataset.

Similarly, the percentage of kept connections after the identification of aperiodicity-unbiased oscillations was higher in the alpha band for all datasets. Connections in the alpha band were preserved by about half after the selection of aperiodicity-unbiased oscillations: in dataset A - HC, 50 % of connections (sd = 21.5) were kept, 44.7 % in dataset A - PD and an average of 77 % (sd = 28.1) for the dataset B. In the beta band, the average percentage of connections did not exceed 10% : an average of 3.1 % (sd = 2.7) and 4.7 % (sd = 6.4) of connections were kept in dataset A-HC and A-PD respectively. In dataset B, about 7.7 % (sd = 14.6) of connections remained. In delta, theta, and gamma bands, the percentage of preserved connections was less than 1% across all datasets.

### Consistency of functional networks computed on aperiodicity-unbiased oscillations across subjects

In order to evaluate if FC between ROIs is consistent between subjects, we computed matrices encoding the percentage of participants exhibiting each edge for each frequency and dataset (Figure 4).

**Figure 4.**
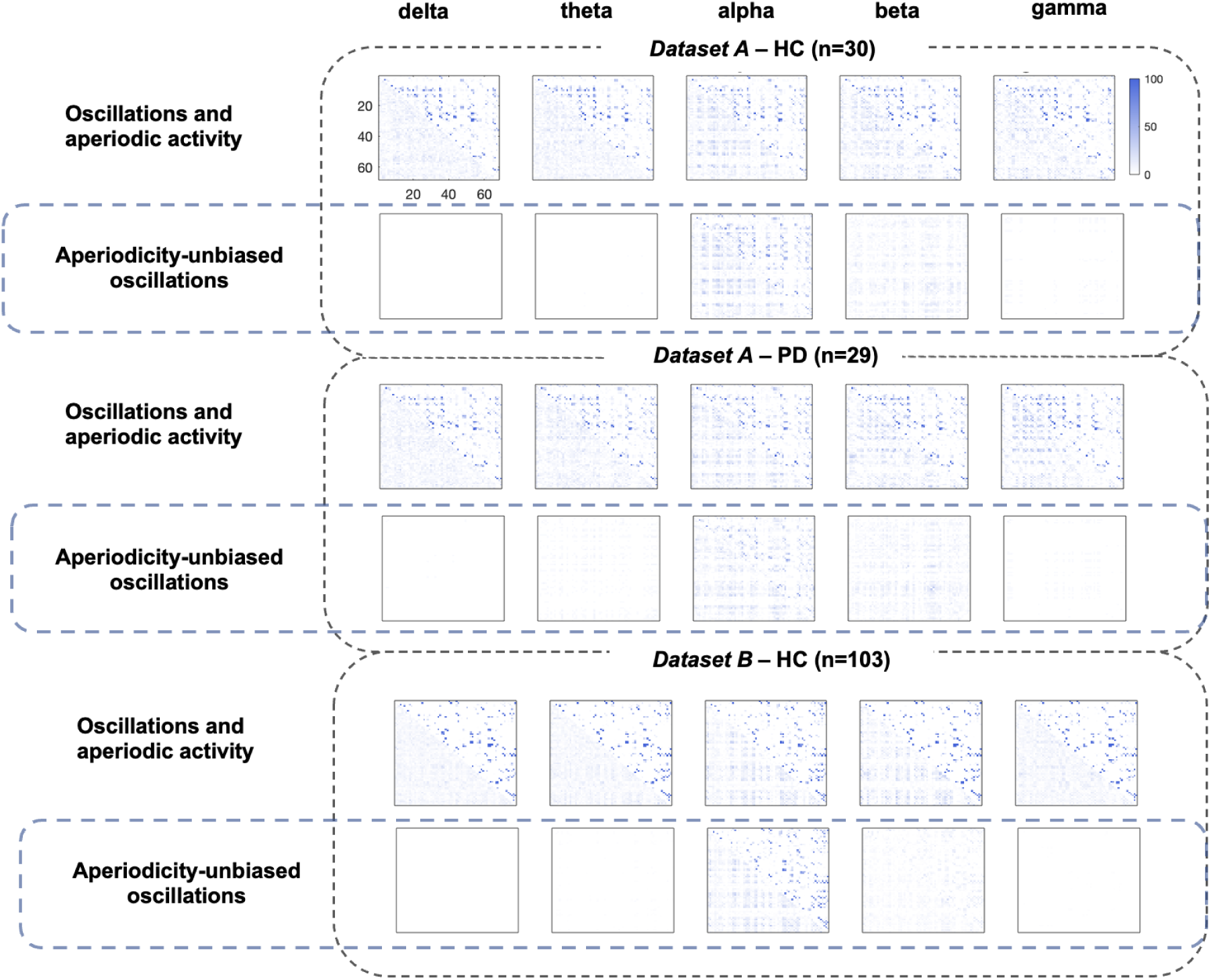
Matrices encoding the percentage of participants exhibiting functional connectivity between regions of interest for each dataset and for each frequency band. The upper triangular portion of the matrix shows the FC results from the PLV method, and the wPLI method for the lower portion. The results are presented for each dataset: the first row presents matrices from the classical pipeline and the second row displays matrices from the approach verifying the presence of oscillations not conflated by aperiodic activity.

First, in the classical pipeline, certain functional connections between ROIs were sometimes present in 100% of the population across all frequency bands. This result was found in all datasets. In our approach, the percentage of participants with edges dramatically decreased for all FC methods and across all frequency bands but the alpha band. More precisely, in our approach, for all datasets and for all FC methods, no participants exhibited FC between ROIs in the delta band. For the alpha band, results were consistent across approaches although the overall percentage of participants showing each connection was reduced. Additionally, the percentage of participants exhibiting FC was strongly reduced in the beta band, with some connections persisting. Results in theta and gamma bands varied across datasets. In the two datasets of healthy participants, no participants exhibited FC in theta band, while 8 PD patients had FC in theta band (x = 46 connexions, sd = 45.5) Some participants in the dataset A showed FC between ROIs in the gamma band, but remaining very sparse, while no participant exhibited FC in this frequency band in the dataset B.

Next, we examined further results in the alpha and beta bands by studying how connections were distributed at the cortical level, and how these were consistent across subjects (Figure 5).

**Figure 5.**
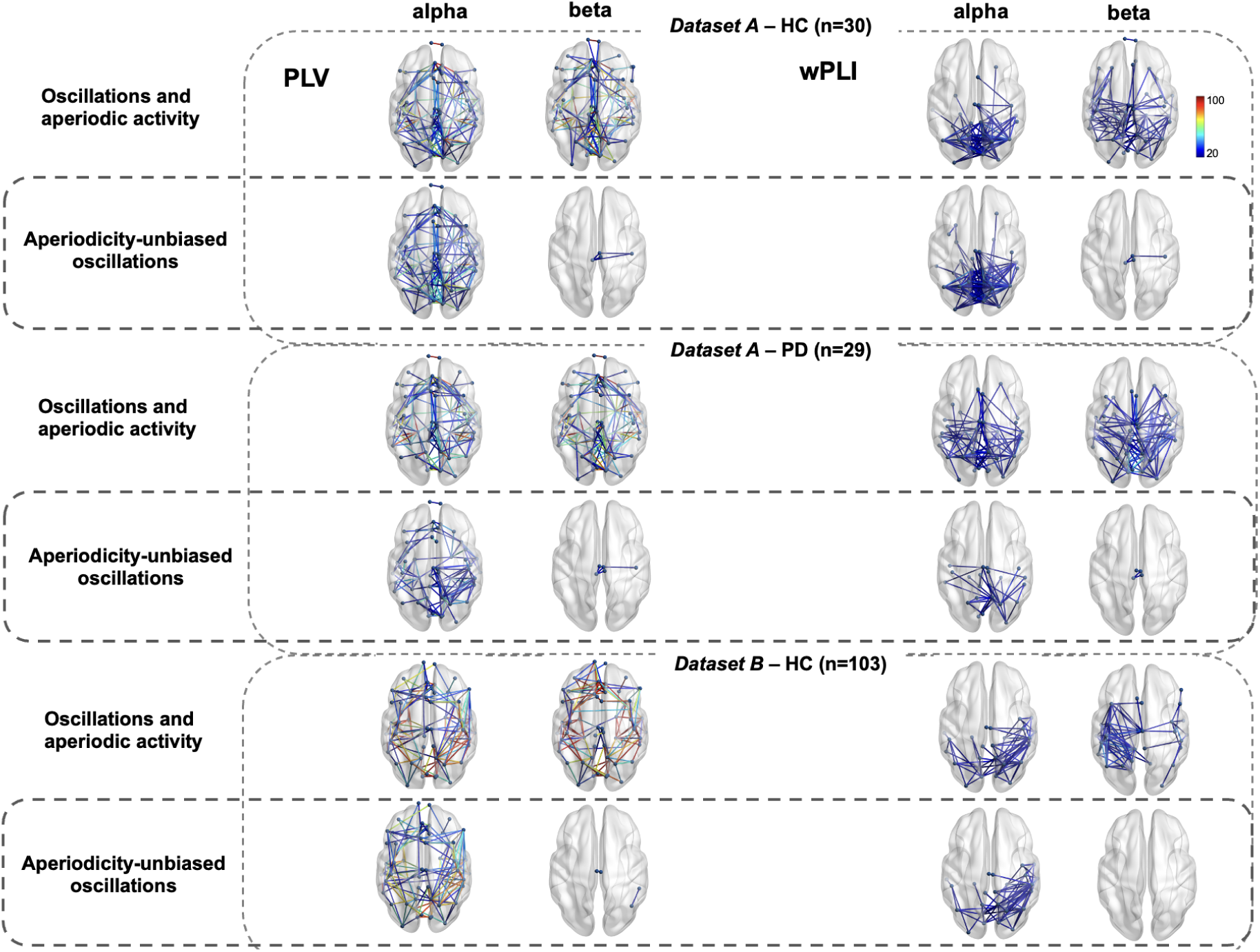
Cortical functional networks in alpha and beta bands across subjects of each dataset for PLV and wPLI. Edges coded the percentage of participants exhibiting the connection. For ease of reading, connections presented were shared by at least 20% of participants. Results are presented for each dataset: the first row shows cortical functional networks from FC matrices from the classical pipeline and the second row displays cortical functional networks from FC matrices from the approach verifying the presence of oscillations not conflated by aperiodic activity.

In the alpha band, PLV functional networks derived from the classical pipeline were diffuse and distributed across the entire cortex. Overall, most connections were shared by less than half of the population, but there were also more widely shared connections, with some even found in the entire population. After selecting the aperiodicity-unbiased oscillations, alpha functional networks exhibited a similar architecture, particularly in healthy subjects, but were shared by fewer individuals.

Regarding the beta band, functional networks were also diffuse across the entire cortex, with some connections shared by the entire population for each dataset. After selecting the aperiodicity-unbiased oscillations, only a few connections, at most four, persisted in less than half of participants. The network was restricted and localized to the medial-parietal region, a result that was consistent across all datasets.

Regarding wPLI functional networks derived from the classical pipeline, a parieto-occipital alpha network was observed, shared by less than half of participants in each dataset. However, PD patients appeared to have a more diffuse network compared to healthy participants. Alpha networks computed on the aperiodicity-unbiased oscillations were more restricted in PD patients, but maintained a parieto-occipital architecture across all datasets. These networks were also shared by less than half of the population. In the beta band, similarly to PLV, networks derived from the classical pipeline were rather diffuse and shared by only few participants, while networks computed on aperiodicity-unbiased oscillations were restricted to a few connections, at most three, localized to the medial-parietal region. These connections were shared by a minority of participants in dataset A, while they were not shared by at least 20% of the population in dataset B.

### Graph theory metrics

In the classical approach, graph metrics had similar values across each frequency band (Figure 6). In our approach, graph metrics values were significantly reduced in corrected networks for all frequency bands and for all datasets. Values were generally null or close to zero in delta, theta, and gamma bands, while a significant decrease was observed in beta and alpha bands (Table 1).

**Figure 6.**
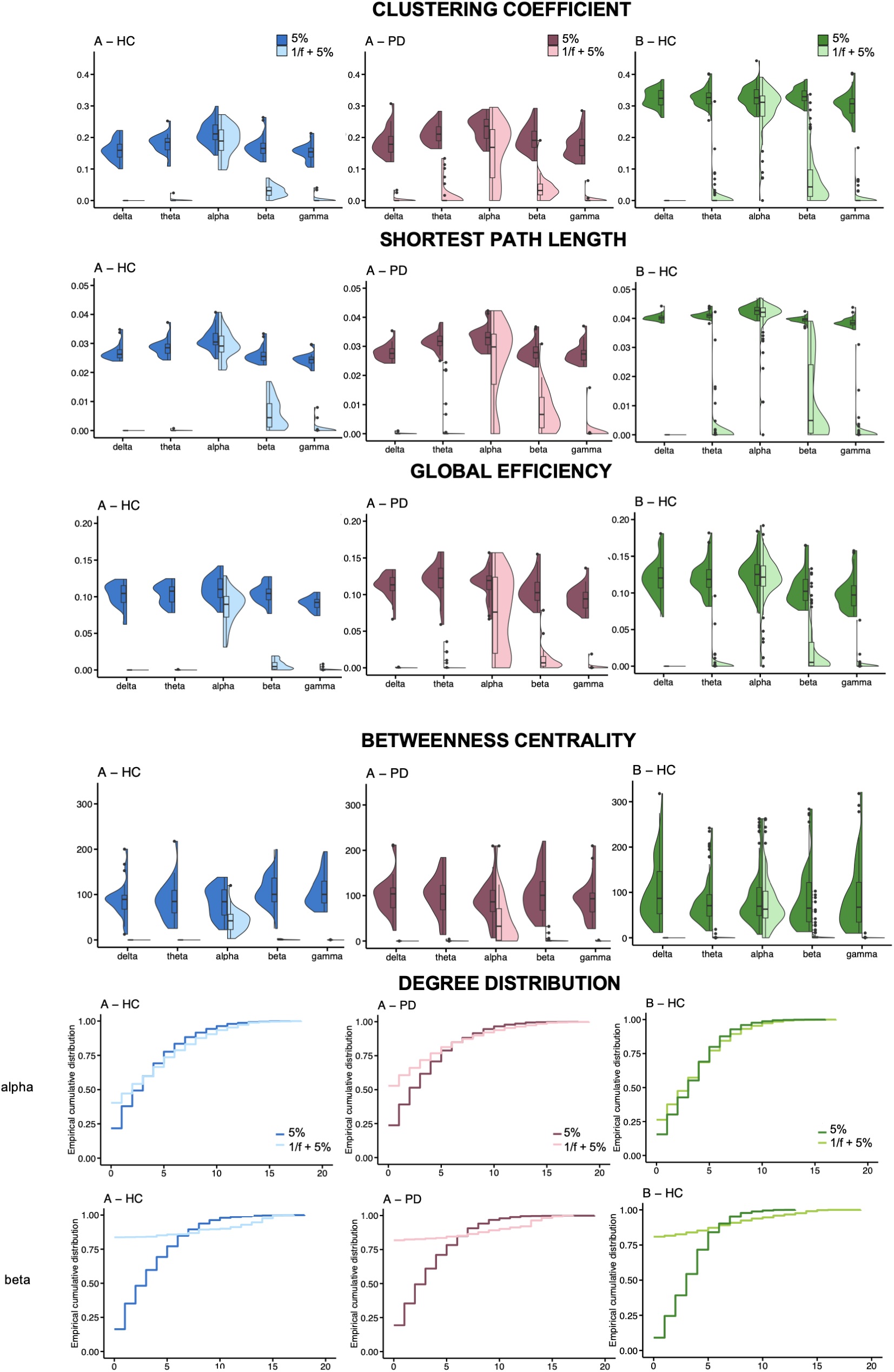
Graph theory metrics of FC networks estimated with PLV according to the approach. The lightest color corresponds to values from matrices computed on aperiodicity-unbiased oscillations while the darkest color indicates values from FC matrices derived from the classical pipeline. The significance of the statistical tests was verified by excluding the outliers. For exact values of statistical tests, please refer to Table 1.

**Table 1.**
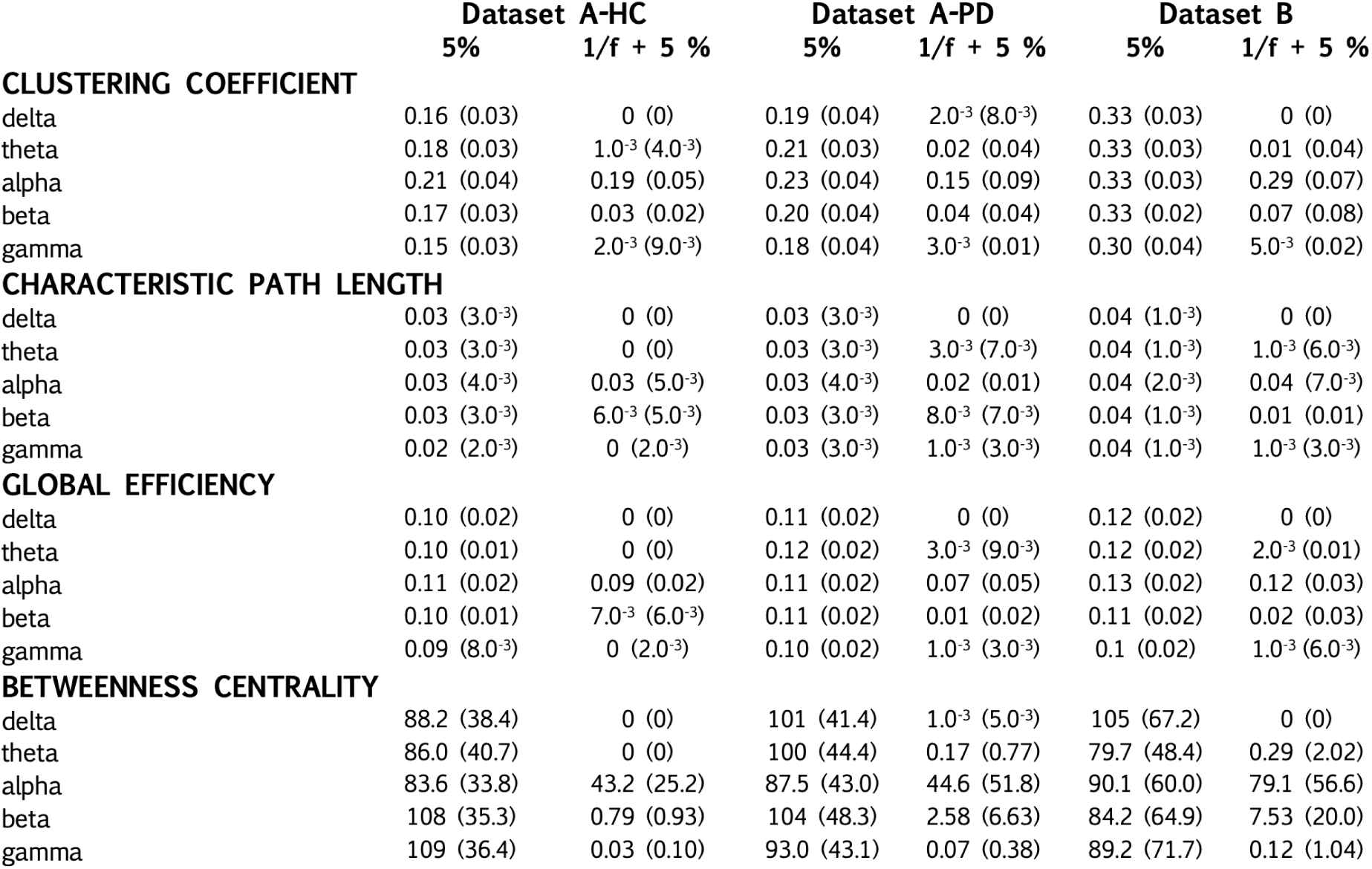
Mean (standard deviation) graph metric values according to the approach for each frequency band and for each dataset. . ‘5%’ refers to the results from the classical pipeline and ‘1/f + 5%’ corresponds to the results from our approach.

First, a significant reduction of clustering coefficient values in networks based on aperiodicity-unbiased oscillations for all frequency bands and all datasets was found. Clustering coefficient values are markedly different for the delta, theta, and gamma bands in all datasets. More particularly, in the alpha band, clustering coefficient are significantly lower after selection of aperiodicity-unbiased oscillations (dataset A: F(1,57)=33.63, p <0.0001; dataset B: t = -5.15, p <0.0001) as well as in the beta band (dataset A: F(1,57) = 819.67, p <0.0001; dataset B: t = -24.54, p <0.0001). The clustering coefficient being a measure of segregation, this suggests that these networks were less segregated.

Similarly, a significant reduction of characteristic path length values was found after the selection of aperiodicity-unbiased oscillations for all frequency bands and in each dataset. Again, differences in path values are marked in the delta, theta, and gamma bands. Characteristic path length values are also significantly reduced with our approach in the alpha band (dataset A: F(1,57)=23.66, p<0.0001; dataset B: t=-3.68,p = 0.0005) and in the beta band (dataset A: F(1,57)= 643.37, p < 0.0001); dataset B: t=-17.90, p < 0.0001). This reduction implies integration between local clusters after selection of aperiodicity-unbiased oscillations.

Global efficiency values were also lower in networks derived from our approach in delta, theta, beta (dataset A : F(1,57)=1474, p <0.0001, dataset B: t = -20.19, p <0.0001) and gamma bands. In the alpha band, global efficiency was reduced in our approach in dataset A (F(1,57)=53.16, p <0.0001) but not in dataset B when outliers were not included (t=-1.32, p = 0.19). These results suggest that networks exhibited less communication efficiency when aperiodicity-unbiased oscillations were selected in all frequency bands for dataset A and in all frequency bands except in the alpha band for dataset B.

Betweenness centrality values were also diminished in our approach in all frequency bands and in each dataset. Differences according to the approach were marked in the delta, theta, beta and gamma bands. Regarding the alpha band, lower values were found after the aperiodicity-unbiased oscillations selection (dataset A: F(1,57)=71.86, p <0.0001; dataset B: t=-2.6, p= 0.01). This significant reduction in betweenness centrality values implies that networks computed on aperiodicity-unbiased oscillations had fewer centrality.

### Consistency of functional connectivity in the alpha band across approaches

For all datasets, alpha functional networks derived from the classical pipeline were highly correlated with alpha functional networks computed on aperiodicity-unbiased oscillations. (Table 2).

**Table 2.** Spearman correlations between alpha functional networks from both approaches for all datasets.

|  |  | **Dataset A-HC** | **Dataset A-PD** | **Dataset B** |
| --- | --- | --- | --- | --- |
| **PLV** | R | 0.95 | 0.95 | 0.99 |
|  | p | 0 | 0 | 0 |
| **wPLI** | R | 0.93 | 0.83 | 0.98 |
|  | p | 0 | 0 | 0 |
| **ciPLV** | R | 0.93 | 0.83 | 0.98 |
|  | p | 0 | 0 | 0 |
| **orthoAEC** | R | 1 | 1 | 1 |
|  | p | 0 | 0 | 0 |
| **AEC** | R | 1 | 1 | 1 |
|  | p | 0 | 0 | 0 |
| **Mean (sd)** |  | 0.96 (0.04) | 0.92 (0.09) | 0.99 (0.01) |

## DISCUSSION

The aim of this study was to evaluate the quantitative contribution of aperiodic activity in the estimation of functional networks. A key finding emerged from our results: aperiodic activity is a major contributor of functional networks in resting state identified by classical methods. We found that EEG oscillatory networks were drastically sparser than classically estimated: >99% of delta, theta, and gamma functional networks; >90% of beta functional networks; and 23-55% of alpha functional networks were actually driven by aperiodic activity and not by oscillatory activity. The number of nodes and percentage of kept connections were greatly reduced in all frequency bands, but remained the highest in the alpha band. Only alpha band activity maintained high FC values, and functional networks in this band were consistent across approaches.

### Overestimation of delta, theta, beta and gamma functional networks in resting state

In delta, theta and gamma bands; accounting for aperiodic activity resulted in dramatically sparser FC matrices. Additionally, both the number of nodes and percentage of preserved connections were drastically reduced, approaching zero. In comparison to the traditional approach where connections were often observed in these frequency bands across the vast majority of individuals, the approach considering only aperiodicity-unbiased oscillations showed a significantly reduced number of connections. In these frequency bands, oscillatory EEG networks were drastically sparser after accounting for aperiodicity-unbiased oscillations, with less than 1% of identified delta, theta, and gamma networks. Our approach resulted in sparser connectivity matrices in the beta band. The percentage of participants exhibiting FC was strongly reduced, with some connections and nodes remaining. While the classical approach showed diffuse functional networks with some connections shared by the entire population, only few connections persisted after accounting for aperiodic activity. Therefore, our results call into question the relevance of considering FC in these frequency bands during resting-state as biomarkers (de Aguiar Neto & Rosa, 2019; Mussigmann et al., 2022) and highlight the need for caution in properly isolating aperiodic activity to study networks based on aperiodicity-unbiased oscillations.

### Consistency of alpha functional network in resting-state

Only FC values in the alpha band were similar between the two approaches. When identifying aperiodicity-unbiased oscillations, connectivity matrices were comparable and correlated with those obtained with the classical approach. The number of nodes and percentage of connections were higher than in other frequency bands. However, it is important to note that the percentage of participants showing connections decreased when considering aperiodicity-unbiased oscillations. Previous reports have indeed evidenced that alpha rhythm dominates brain activity at rest (Samogin et al., 2020; Roopun et al., 2008) and that alpha networks are linked to the default network (Mantini et al., 2007).

### On the importance of a new methodological “good practice”: verifying the presence of oscillations before estimating functional connectivity

Our analyses highlight the importance of establishing a new methodological good practice: verifying the presence of aperiodicity-unbiased oscillations in ROIs before proceeding to the estimation of the correlation between these oscillations, which defines FC. Indeed, rigorous scientific practice requires treating the presence of true oscillations as a fundamental assumption when calculating FC, since it is intended to quantify communication between ROIs based on the theory that oscillatory synchronization facilitates communication. Consistent with previous studies, our findings evidence that aperiodic activity biases the measurement of putative oscillations, and highlight the importance of parameterizing the neural power spectrum as a combination of aperiodic and periodic activity for accurate interpretation of the EEG signal (Del Bianco et al., 2024; Donoghue et al., 2020; Donoghue & Watrous, 2023; Ostlund et al. 2022).

### Methodological limitations

The *specparam* algorithm has some methodological limitations, but its spectral parameterization remains at least as performant as other methods such as BOSC or IRASA (Donoghue et al., 2020). A recent study also evidenced that the test-retest reliability of parameterized periodic and aperiodic activity with *specparam* is satisfactory in eyes-closed resting state (Mc Keown et al., 2024). Differences between the datasets studied, such as sample size, number of cap electrodes, age population should be taken into consideration, but the replication of our results despite these differences strengthens our findings’ robustness. The same applies to preprocessing differences; despite the various pipelines between the two datasets, results were nearly identical in showing the dramatically sparse nature of FC networks at rest (except for the alpha band), further supporting the robustness of our findings. Additionally, another limitation is that an implicit assumption of these analyses is that oscillations are considered to be sinusoidal. It may be interesting to investigate whether this assumption holds true when considering more complex and realistic waveforms for electrophysiological signals.

### Conclusions and perspectives

Given that this study focused on resting-state, it would be important to further investigate the impact of considering aperiodic activity on the estimation of functional networks associated with the execution of cognitive tasks, which could be differentially associated with oscillatory activity. Our results strongly suggest that considering aperiodicity-unbiased oscillations is a necessary prerequisite to estimate functional networks when one wants to rely on the assumption that neuronal communication is facilitated by oscillatory synchronization.

## ACKNOWLEDGMENTS

We warmly thank all the volunteers who participated in this study.

