## Supplementary Material for "Functional connectivity is dominated by aperiodic, rather than oscillatory, coupling"

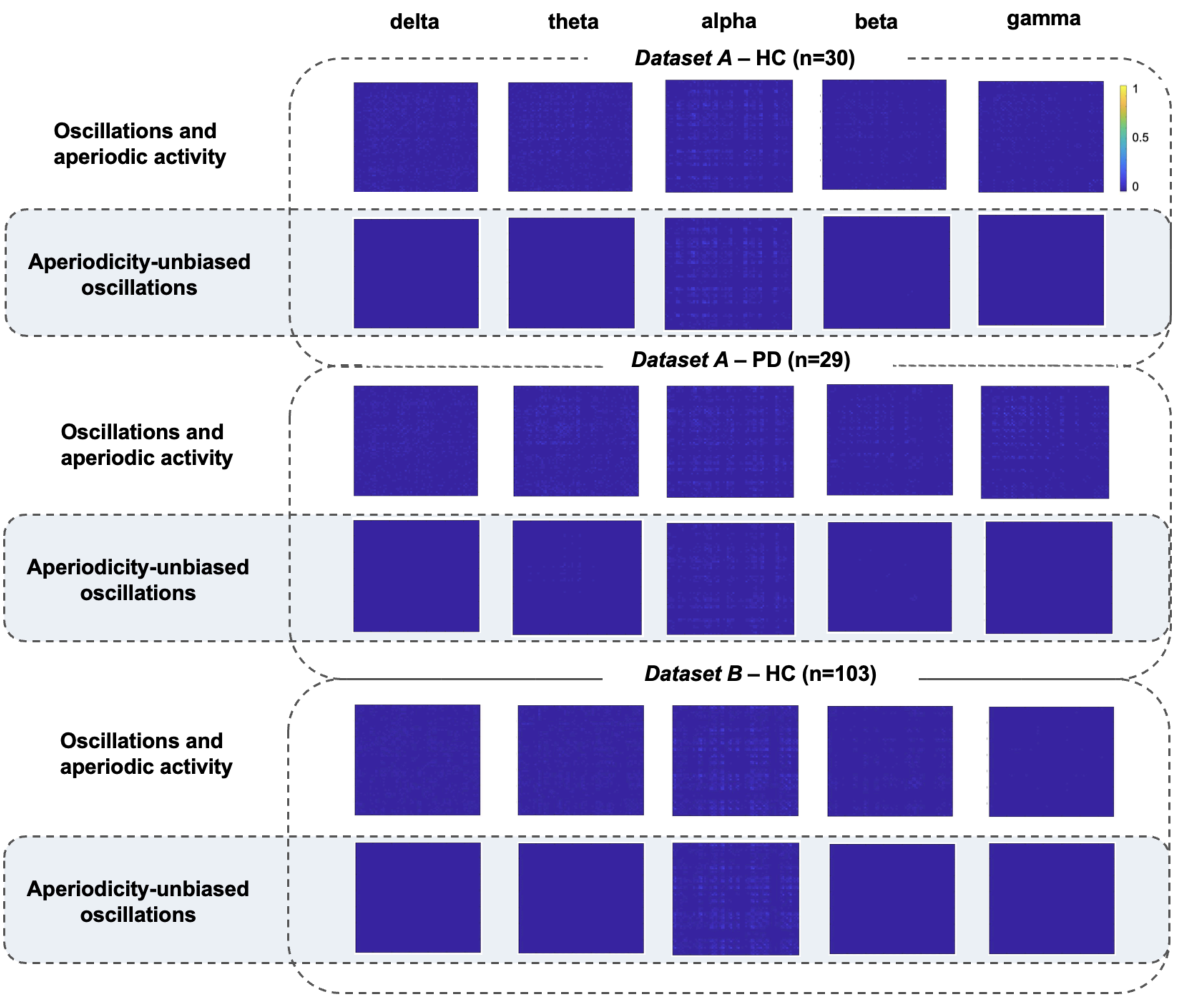


**Supplementary Figure 1 Functional connectivity matrices, using ciPLV, averaged across subjects for each dataset and each frequency band**. Results are presented for each dataset: the first row shows FC matrices from the classical pipeline and the second row displays FC matrices from the approach verifying the presence of oscillations not conflated by aperiodic activity.

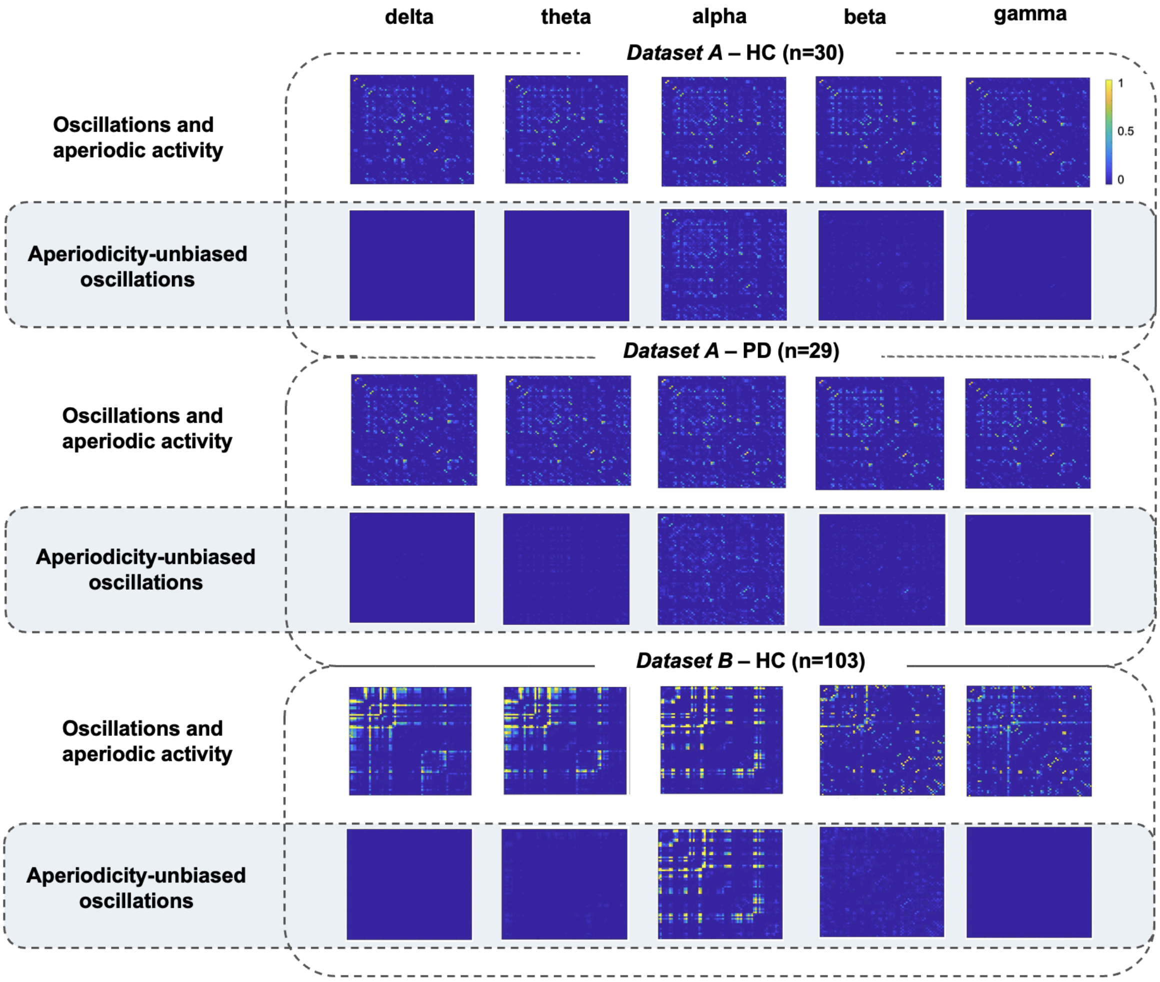


**Supplementary Figure 2 Functional connectivity matrices, using AEC, averaged across subjects for each dataset and each frequency band**. Results are presented for each dataset: the first row shows FC matrices from the classical pipeline and the second row displays FC matrices from the approach verifying the presence of oscillations not conflated by aperiodic activity.

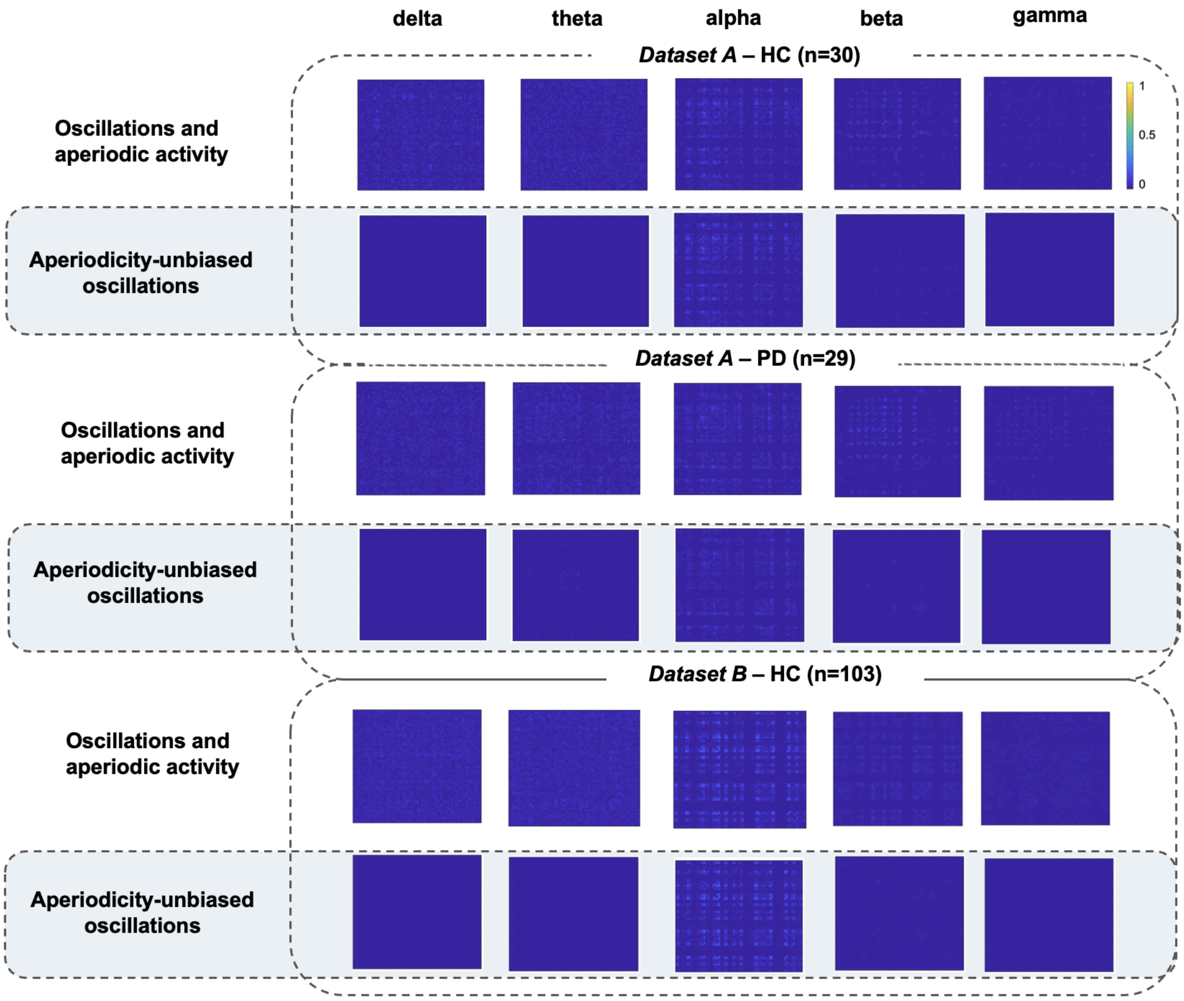


**Supplementary Figure 3 Functional connectivity matrices, using orthoAEC, averaged across subjects for each dataset and each frequency band**. Results are presented for each dataset: the first row shows FC matrices from the classical pipeline and the second row displays FC matrices from the approach verifying the presence of oscillations not conflated by aperiodic activity.
